# Mechanical control of tissue shape and morphogenetic flows during vertebrate body axis elongation

**DOI:** 10.1101/2020.06.17.157586

**Authors:** Samhita P. Banavar, Emmet K. Carn, Payam Rowghanian, Georgina Stooke-Vaughan, Sangwoo Kim, Otger Campàs

## Abstract

Shaping embryonic tissues into their functional morphologies requires cells to control the physical state of the tissue in space and time. While regional variations in cellular forces or cell proliferation have been typically assumed to be the main physical factors controlling tissue morphogenesis, recent experiments have revealed that spatial variations in the tissue physical (fluid/solid) state play a key role in shaping embryonic tissues. Here we theoretically study how the regional control of fluid and solid tissue states guides morphogenetic flows to shape the extending vertebrate body axis. Our results show that both the existence of a fluid-to-solid tissue transition along the anteroposterior axis and the tissue surface tension determine the shape of the tissue and its ability to elongate unidirectionally, with large tissue tensions preventing unidirectional elongation and promoting blob-like tissue expansions. We predict both the tissue morphogenetic flows and stresses that enable unidirectional axis elongation. Our results show the existence of a sharp transition in the structure of morphogenetic flows, from a flow with no vortices to a flow with two counter-rotating vortices, caused by a transition in the number and location of topological defects in the flow field. Finally, comparing the theoretical predictions to quantitative measurements of both tissue flows and shape during zebrafish body axis elongation, we show that the observed morphogenetic events can be explained by the mere existence of a fluid-to-solid tissue transition along the anteroposterior axis. These results highlight the role of spatiotemporally-controlled fluid-to-solid transitions in the tissue state as a physical mechanism of embryonic morphogenesis.

## Introduction

During embryonic development, tissues undergo major physical transformations to build functional structures. Similar to inert materials, shaping embryonic tissues necessarily involves the spatiotemporal control of several key physical quantities^1^, namely its growth (e.g., cell proliferation), material properties and/or active stresses. However, unlike inert systems, living tissues are active materials and can locally regulate the value of these fields through local changes in cell behavior. In general, it is unclear what physical fields are spatiotemporally controlled to sculpt tissues and organs, mainly because measurements of spatiotemporal variations in these physical quantities within developing embryos are still sparse and challenging. Since spatiotemporal variations in multiple physical fields can contribute to the morphogenetic processes^1^, it is important to have information on all these fields in the same system to establish how the tissues are physically shaped.

Recently, quantitative measurements of the spatial variations in both mechanical stresses and tissue material properties showed the existence of a fluid-to-solid transition in the state of the tissue during the posterior extension of the body axis in zebrafish embryos^2^ (Fig. 1a). During zebrafish posterior body elongation, cells in dorsal-medial (DM) tissues continuously move ventrally to the mesodermal progenitor zone (MPZ) (Fig. 1b,c), providing the necessary material to extend the body axis since, at the developmental stages studied, cell proliferation is negligible and does not substantially contribute to the elongation of the body axis^3, 4^. The mesodermal progenitor cells in the MPZ progressively incorporate into the presomitic mesoderm (PSM), a process that involves their maturation into mesodermal cells and their gradual arrest^2, 5–7^. The observed fluid-to-solid transition was found to be associated with cellular jamming along the anteroposterior (AP) axis following an anterior decrease in extracellular spaces and in cell-cell contact active fluctuations^2^. Regardless of the specific physical mechanism of this transition, the tissue was found to switch from a fluid-like state in the posterior end of the body, namely the MPZ, to a solid-like state in the PSM (Fig. 1a-c). Continuum mechanics simulations showed that the existence of a fluid-to-solid transition in the tissue could reproduce the observed unidirectional body axis elongation^2^, but it remains unclear what types of morphogenetic flows and tissue shapes (morphological phenotypes) can be achieved with the observed transition, how different physical parameters control morphogenetic changes, and how different physical fields (stress, tissue pressure, velocity, etc.) vary spatially during body axis elongation. Our goal here is to answer these questions by performing extensive simulations of the process and comparing the results to quantitative measurements of tissue flows and shape during zebrafish posterior axis elongation.

**Figure 1.**
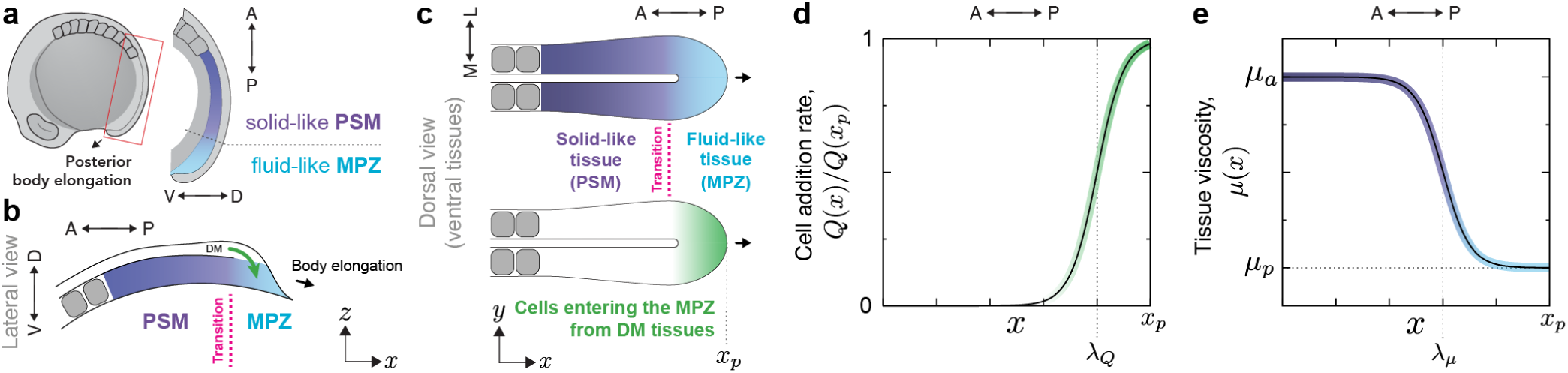
Fluid-to-solid tissue transition during zebrafish posterior axis elongation. **a**, Schematic lateral view of the zebrafish embryo at 10-somite stage, showing the previously reported^2^ fluid-like mesodermal progenitor zone (MPZ) and solid-like presomitic mesoderm (PSM). **b**, Cells from dorsal-medial (DM) tissues (green arrow) enter the ventral MPZ tissue at the posterior end (lateral view). The tissue undergoes a fluid-to-solid transition along the AP axis, as cells from the fluid-like MPZ mature and join the solid-like PSM. **c**, Sketch of a dorsal view of ventral tissues showing both the fluid-to-solid tissue transition and the entrance of cells from the DM into the MPZ. **d-e**, Spatial profiles of the rate of cell (material) addition to ventral tissues, *Q*(*x*) (d), and tissue viscosity, *μ*(*x*) (e), used in the simulations.

Several methods exist to simulate tissue morphogenesis^8, 9^. Cell-based models are well-suited when cellular resolution is necessary, but typically involve a large number of parameters (or assumptions of such parameters values) because the mechanical state of each cell needs to be specified. Moreover, since the tissue material properties and stresses emerge from the collective behavior of cells, the connection between mechanical parameters at the cell scale and material properties at the tissue scale can be quite complex. When studying tissue morphogenesis at length and time scales characteristic of tissue dynamics (larger than those of cell dynamics), coarse-grained continuum approaches that only require information of physical fields at supracellular scales are better suited^8, 10^. Previous continuum descriptions of tissue morphogenesis generally assumed spatially uniform mechanical properties (i.e., constant tissue viscosity or constant stiffness depending on the tissue) and considered only spatial variations in either cell proliferation or active forces because experimental studies had been mostly focused on these quantities^11–14^. The role of spatial variations in tissue mechanical properties and, especially, the role of regional changes in fluid and solid tissue states, to control embryonic morphogenesis remains largely unexplored.

In the specific case of body axis elongation, self-propelled particle descriptions have been used to understand cellular movements in the tissue^7^. These simulations assumed a set tissue shape (fixed tissue boundaries), allowing the prediction of cell movements but not of tissue morphogenesis since the boundaries were, by construction, fixed. Importantly, most cases of tissue morphogenesis are examples of so-called free boundary problems, in which tissue flows change the shape of the tissue and these boundary changes affect, in turn, morphogenetic flows. It is thus valuable to consider the coupled dynamics of the tissue shape (boundary) and morphogenetic flows when studying theoretically embryonic morphogenesis.

Building on previous work^2^, we theoretically explore the role of a spatially-controlled fluid-to-solid tissue transition on the morphogenetic events that guide the elongation of the zebrafish body axis. We treat the system as a free-boundary problem and perform 2D finite element numerical simulations of tissue morphogenesis based solely on first principles (mass and momentum balance). Our results show that the mere presence of a fluid-to-solid transition along the AP axis enables unidirectional tissue elongation. In the absence of the fluid-to-solid transition, the tissue expands isotropically when the tissue surface tension is sufficiently large. For unidirectional axis elongation, our results show the existence of a sharp transition in the structure of morphogenetic flows without any qualitative change in the underlying tissue mechanics. For small tissue surface tensions, tissue flows display counter-rotating vortices as the tissue transits from fluid-like to solid-like states, whereas at large tissue surface tensions, tissue flows smoothly passage from posterior-directed movements in the MPZ to anterior-directed tissue flow in the PSM. The predicted AP axial stresses indicate that the MPZ tissues push the body posteriorly, contributing to axis elongation, whereas PSM tissues are compressed, indicating that the PSM mechanically sustains the extension of the body. Finally, our predicted tissue flows and shape for small tissue surface tension quantitatively agree with measurements of morphogenetic flows and tissue shape during zebrafish axis elongation.

## Theoretical description

Since we are interested in tissue morphogenesis at supracellular length scales and developmental time scales, we describe the tissue as a coarse-grained continuum. Indeed, all observed mechanical gradients in the tissue during body axis elongation occur at length scales much larger than the cell size and are persistent over timescales longer that characteristic timescales of cellular processes^2^, indicating that a coarse-grained description is apt, as previously done in other systems^8, 10^. Moreover, since the ventral tissues (including MPZ and PSM) are thin along the dorsal-ventral axis (DV, *z* axis) compared to its medial-lateral (ML, *y* axis) and anterior-posterior (AP, *x* axis) extensions^2, 6^ (Fig. 1a,b), we approximate ventral tissues as a 2D system, neglecting the DV tissue thickness, and simulate a 2D DV projection of the ventral tissues (Fig. 1b,c). Finally, since the notochord has little ML extent and the elongation of the body axis has been shown to proceed in its absence^15^, we neglect it here.

In order to sustain the continuous posterior extension of the body axis, it is necessary to constantly add new (tissue) material at the posterior end of ventral tissues. Cell proliferation could potentially contribute to the addition of new tissue material, but it has been previously shown that proliferation is minimal in the tissue and that its inhibition does not preclude the formation of the body axis^4, 16^, indicating that cell proliferation is not driving tissue elongation in zebrafish at these developmental stages^3^. From the perspective of ventral tissues, the dorsal to ventral movement (ingression) of cells at the posterior end of the tissue represents an addition of material to the MPZ region as body elongation proceeds (Fig. 1b,c). However, cell ingression into the MPZ can only occur if enough space can be made available for the ingressing cells, which depends directly on the local tissue pressure since local volume changes are directly coupled to the local value of the pressure: large tissue pressure in the MPZ will prevent ingression of cells from DM tissues because these cannot generate enough force to push their way into the MPZ. Since the ingression of cells to the MPZ from DM tissues is spatially restricted to a region of limited size at the extending posterior-most part of the tissue^7^, we define the rate of cell ingression from DM tissues into the MPZ, *Q*(*x, P*), with both explicit spatial and pressure dependences, namely

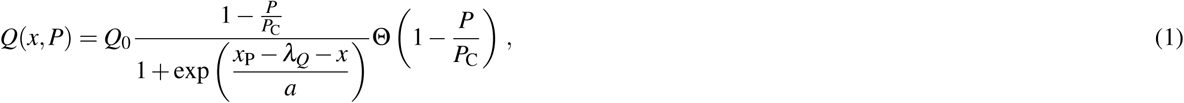

where *P* is the tissue pressure, *P*_*C*_ is the critical pressure over which cells cannot ingress into the MPZ and *Q*_0_ is the maximal cell addition rate at the posterior-most end of the body axis for negligible tissue pressures. The function Θ(•) represents the Heaviside step function and *x*_p_ is the time-dependent position of the end of the body axis. In the case that tissue pressure is small compared to *P*_*c*_, the cell addition rate *Q*(*x, P*) decays along the AP axis from a maximal value *Q*_0_ at the posterior-most end to vanishing values at length scales larger than *λ*_*Q*_, which sets the size of the ingression region (Fig. 1b-d). In this case, *Q* changes from *Q*_0_ to zero over a spatial range of size *a* (with *a ≪ λ*_*Q*_). If the tissue pressure is not negligible compared to *P*_*c*_, cell ingression will be limited further according to the spatial profile of the tissue pressure, and eventually halted for tissue pressures above *P*_*c*_. In the reference frame of the extending posterior end, the profile *Q* does not change over time (Fig. 1d), but in the absolute reference frame it does so through its dependence on the position *x*_p_(*t*) of the extending body end.

In order to simulate the physical growth of the tissue, it is necessary to know the spatiotemporal changes in tissue material properties. As explained above, direct measurements of tissue mechanics have revealed that ventral tissues undergo a fluid-to-solid transition along the AP axis, with the fluid-like MPZ tissues rigidifying into to a solid-like PSM^2^. This transition was associated to cellular jamming, in the broad sense of the word within the classification of jamming transitions^17^. More specifically, the observations resemble more closely a glass transition, as actively-generated cell-scale forces were shown to create cell-cell contact fluctuations that can be qualitatively thought of as an effective temperature. In glass transitions, the viscosity of the system becomes arbitrarily large as the system is cooled below the glass transition temperature^18^, becoming effectively a solid. Within this framework, the fluid-like state of the extending posterior MPZ tissue can be qualitatively thought of as the tissue having an effective temperature higher than the glass transition temperature, enabling cellular rearrangements and tissue fluidization. In contrast, the solid-like PSM can be thought of as a tissue with an effective temperature lower than the glass transition temperature, leading to very large viscosities that barely allow any tissue reorganization at the observation timescales, effectively rigidifying the PSM. To account for a fluid-to-solid transition of this nature along the AP axis, we describe the tissue as a viscous fluid with inhomogeneous viscosity *μ*(*x*), minimal at the posterior end of the body and sharply, but smoothly, transiting to a high viscosity value at a defined distance *λ*_*μ*_ from the posterior end of the body (Fig. 1b,e), namely

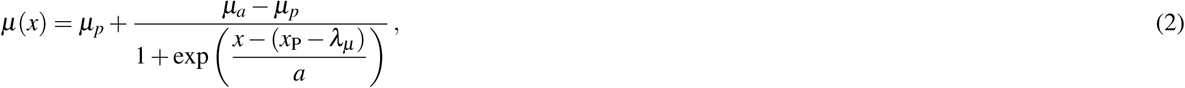

where *μ*_*p*_ and *μ*_*a*_ are the values of the tissue viscosity in the MPZ and PSM, respectively. The transition between low and high tissue viscosities occurs over a region of size *a* (with *a ≪ λ*_*μ*_). Large values of *μ*_*a*_*/μ*_*p*_ (*μ*_*a*_*/μ*_*p*_ *→* ∞) simulate the observed fluid-to-solid tissue transition, but it is also possible to simulate a tissue with uniform viscosity (*μ*_*p*_ = *μ*_*a*_) and intermediate behaviors.

Knowing the spatial distribution of the cell ingression rate and tissue viscosity along the AP axis, which we consider here as input fields, it is possible to simulate the dynamics of tissue morphogenesis. Two fundamental equations govern the dynamics of the system, namely mass conservation (or mass balance) and momentum conservation. In the presence of spatially-dependent cell ingression, Q(*x, P*), mass conservation reads

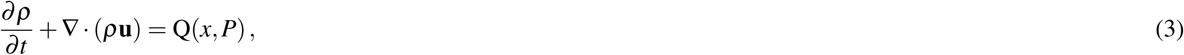

where **u** and *ρ* are the velocity and density fields, respectively. Since the cell density has been experimentally shown to be uniform along the AP axis^2^, we assume *ρ* to be constant in what follows, which reduces Eq. 3 to

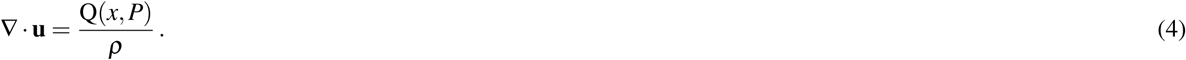

At the typical length scales involved (∼100 *μ*m) and for the measured values of tissue viscosity (∼10^5^ Pa s^2, 19^), the dynamics can be safely assumed to be overdamped. In these conditions, momentum conservation reduces to force balance, which reads

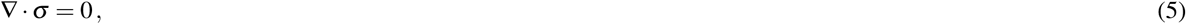

where *σ* is the stress tensor. For a viscous fluid with inhomogeneous viscosity *μ*(*x*) in 2D, the stress tensor reads

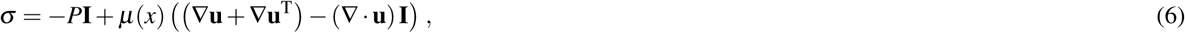

where **I** is the identity tensor. This pressure does not correspond to any hydrostatic pressure in the tissue, but rather is the crowding pressure between cells in the tissue, mirroring the osmotic pressure in an aqueous foam^20^. While the density in the tissue is constant, the divergence of the velocity field does not generally vanish in Eq. 6 because of the addition of new material (see Eq. 4). The tissue is assumed to be immersed in a fluid environment similar to water (Newtonian fluid) with uniform viscosity several orders of magnitude smaller than that of the tissue. The equations governing the dynamics of the surrounding fluid are also mass and momentum conservation, but in this case, there are no sources of material.

To solve the equations above, it is necessary to specify the boundary conditions. The shape of the tissue is not imposed in any way and depends on the physical fields inside the tissue. In the same way, these physical fields depend on the location of the boundary, i.e., the shape of the tissue. As for free-boundary problems related to the dynamics of fluid-fluid interfaces^21^, the boundary conditions are velocity continuity and local normal force balance at the tissue boundary (or surface). Continuity of the velocity field simply reads

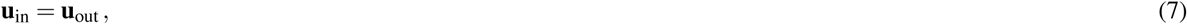

where **u**_in_ and **u**_out_ are the velocities of the tissue and outer fluid surrounding it, respectively, evaluated at the tissue boundary. Local normal force balance (Laplace’s Law) reads

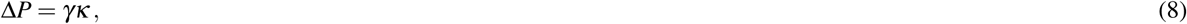

where Δ*P* is the tissue pressure jump at the boundary, *γ* is the tissue surface tension and *κ* is the curvature of the tissue surface. The interface between the tissue and surrounding fluid is described with a single curvature *κ* along its arc-length because the simulations are in 2D. Since the tissue pressure is not associated with any hydrostatic pressure, but is rather a pressure associated with cellular crowding, its value outside the tissue vanishes. The tissue pressure jump Δ*P* at the tissue surface is thus Δ*P* = *P*_*S*_, where *P*_*S*_ is the tissue pressure at the tissue boundary and, in general, varies on the tissue surface. The tissue surface tension *γ* accounts for the surface tension known to exist in multicellular systems with adhering cells^22, 23^. For simplicity, we assume here that at the relevant developmental time scales and supracellular scales the tissue surface tension is spatially uniform and constant in time.

Since the addition of material in the MPZ is essential to sustain body elongation, we scale all lengths with the characteristic length scale *λ*_*Q*_, time with the characteristic timescale of cell ingression, namely *τ ≡ ρ/Q*_0_, and stresses with the critical pressure *P*_*C*_ at which cell ingression to the MPZ ceases. Scaling all variables and equations with the mentioned scales, we obtain the relevant dimensionless parameters that govern the dynamics of the system, namely

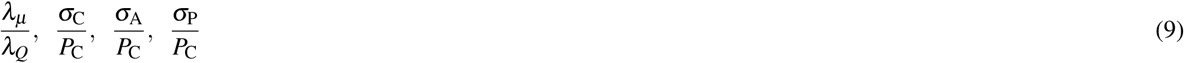

where *λ*_*μ*_ */λ*_*Q*_ is the ratio of the length scale over which tissue viscosity varies to the size of the region where cell ingression occurs (or tissue material is added, equivalently). Beyond this ratio, the other dimensionless parameters are ratios of the four relevant stress scales in the problem, namely the shear stress scales in the MPZ and PSM, *σ*_P_ and *σ*_A_, respectively, the capillary stress associated with tissue surface deformations, *σ*_C_ *≡ γ/λ*_*Q*_, and the critical tissue pressure *P*_C_ over which cell ingression ceases. In analogy to fluid interfaces, the capillary stress measures the necessary stress to deform the tissue surface. The ratio of shear stress scales directly relates to the ratio of tissue viscosities in each region, such that *σ*_P_*/σ*_A_ = *μ*_P_*/μ*_A_, with *μ*_P_*/μ*_A_ = 1 for uniform viscosity and *μ*_P_*/μ*_A_ *→* 0 for a jamming transition at vanishing tissue effective temperature^20^.

In order to narrow the parameter space, we use known experimental values for some parameters. Measurements of the size of the MPZ^2^, *λ*_*μ*_, and the size of the ingression region *λ*_*Q*_ (see below) indicate that *λ*_*μ*_ */λ*_*Q*_ is in the range 1 *< λ*_*μ*_ */λ*_*Q*_ *<* 2. The range of *σ*_P_*/σ*_A_ explored is 10^−3^ − 1, because we are both interested in the limit of uniform tissue viscosity (*μ*_A_ = *μ*_P_) and in the presence of a fluid-to-solid transition (*μ*_A_ ≫*μ*_P_). We considered the ratio of capillary to critical pressure controlling *P*_C_, *σ*_C_*/P*_C_, to vary over a range 0.01 − 10. We checked that the results show negligible dependence on the transition zone size *a* as long as *a* is sufficiently small. Consequently, we fix *a* = *λ*_*Q*_*/*4 in our simulations.

## Results

### Morphogenesis of the posterior vertebrate body axis

To understand the possible tissue shapes and their dynamics, we numerically integrated Eqs. 4-5 and obtained the time evolution of the system for different parameter values (Methods). Starting from an initial semicircular tissue shape (Fig. 2a; Methods), we let the tissue shape evolve over time for different parameters and identify the different dynamical regimes. For values of the capillary stress larger than a threshold value in the critical pressure, namely *σ*_C_*/P*_*C*_ *≃* 4, the tissue cannot extend in any way and remains arrested (Fig. 2a). This is because tissue material (cells) cannot enter the MPZ due to the high crowding pressure in the MPZ (Fig. 2b), thus halting growth (Fig. 2c). This high pressure in the tissue is a direct consequence of the large capillary stress (compared to *P*_*C*_) that resists deformation and extension of the tissue surface.

**Figure 2.**
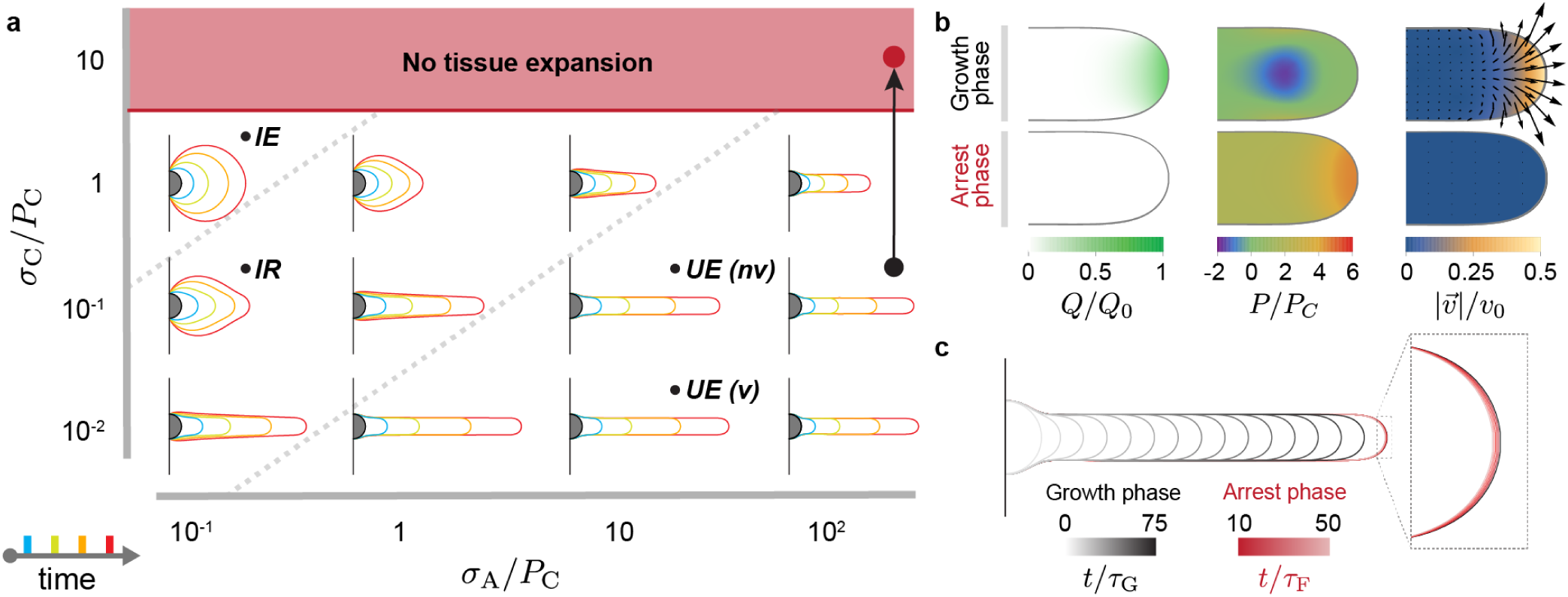
Morphogenesis of posterior tissues. **a**, Time evolution of posterior tissue shape from an initial semi-circular shape (gray), as *σ*_C_*/P*_*C*_ and *σ*_A_*/P*_*C*_ are varied (*σ*_A_*/σ*_P_ = 10, *λ*_*μ*_ */λ*_*Q*_ = 2). Four time snapshots at *t/τ* = 10, 25, 50, 75 (blue, green, orange and red, respectively) are shown for each parameter combination. The dotted lines qualitatively separate different regimes: isotropic expansion (*IE*), intermediate regime (*IR*), and unidirectional elongation (*UE*). **b-c** A sharp increase in capillary stress during unidirectional elongation (arrow in a), leads to an increase of tissue pressure (b) that prevents addition of material from dorsal tissues into the MPZ (b; vanishing *Q*), halting tissue flows (b; vanishing velocities) and arresting further tissue elongation (c).

Below the threshold value causing growth arrest (i.e., *σ*_C_*/P*_*C*_ ≲ 4), the tissue can extend, albeit differently for varying values of parameters. When the capillary stresses are much larger than the shear stresses both in the MPZ and the PSM (*σ*_C_ ≫*σ*_P_ and *σ*_C_ ≫ *σ*_A_), it is much more costly to deform the tissue surface than to induce material flows within it. Consequently, in this regime, the tissue expands isotropically keeping a round shape, as would a liquid drop with high surface tension with liquid being injected in it (Fig. 2a). In contrast, when the capillary stresses are large compared to the shear stresses in the MPZ (*σ*_P_ ≪*σ*_C_), but small compared to the shear stresses in the PSM (*σ*_A_ ≫ *σ*_C_; i.e., if the viscosity *μ*_A_ is large enough), the MPZ tissue can easily flow upon addition of new cells from dorsal tissues, but the PSM can barely flow within the timescales of tissue growth. Since anterior PSM tissues do not flow due to their large viscosity in this limit, they effectively behave as a solid at the timescales relevant to tissue morphogenesis. In this case, it is much less costly to deform the tissue at the posterior end than to induce flows in the PSM and, as a result, the tissue extends unidirectionally and posteriorly (Fig. 2a). This situation, in which the PSM effectively behaves like a solid and the MPZ behaves like a fluid, corresponds to experimentally observed fluid-to-solid tissue transition from MPZ to PSM^2^. Since unidirectional axis elongation can only be achieved when the capillary stresses associated with the tissue surface tension are smaller than the shear stresses in the PSM, our results suggest that capillary stresses associated with the tissue surface tension are small in zebrafish posterior tissues compared to the other stress scales in the system. In between these two limiting regimes (purely isotropic growth and unidirectional elongation), there is an intermediate regime that displays some characteristics of both. If the capillary stress scale becomes comparable to the shear stress scales (*σ*_C_ ∼ *σ*_A_ and *σ*_C_ ∼ *σ*_P_), then the tissue expands mostly isotropically but also displays a posterior bump in the tissue shape due to the localized addition of cells in that region (Fig. 2a). In this case, the viscosity in the PSM is not large enough to support the posterior unidirectional extension of the tissue and prevent mediolateral tissue spreading over the timescales of tissue morphogenesis, but it is not small enough either to fully prevent it. As a consequence, the tissue spreads mediolaterally at the anterior end, leading to a blob-like anterior tissue expansion in this region, while displaying a posteriorly extending bump at the posterior end.

The different dynamical regimes of tissue expansion (Fig. 2a) will, in general, be characterized by different morphogenetic flows. In the case of isotropic tissue expansion with uniform tissue viscosity (*σ*_C_ *> σ*_A_ = *σ*_P_), strong mediolateral and anterior-directed flows are present, which redistribute the tissue material added at the posterior end (Fig. 3a). Indeed, in this regime the capillary stresses are larger than all shear stresses in the tissue, forcing the material added at the posterior-most tissue end to redistribute mediolaterally and anteriorly while preserving a nearly spherical tissue shape during tissue expansion. Since the shear stresses are much smaller than the capillary stress, the tissue can easily flow and quickly reduce pressure differences, leading to an almost uniform tissue pressure inside the isotropically expanding tissue. In the intermediate regime (Fig. 2a and Fig. 3c), the morphogenetic flows show the same characteristics of isotropic growth (Fig. 3a), namely anteriorly-oriented and mediolateral flows. Moreover, the largely uniform pressure is also characteristic of isotropic growth, indicating that while the shape in this intermediate regime displays a posterior bump (because shear stresses in this regime are large enough to start deforming the tissue boundary), the morphogenetic process is qualitatively akin to isotropic tissue expansion.

**Figure 3.**
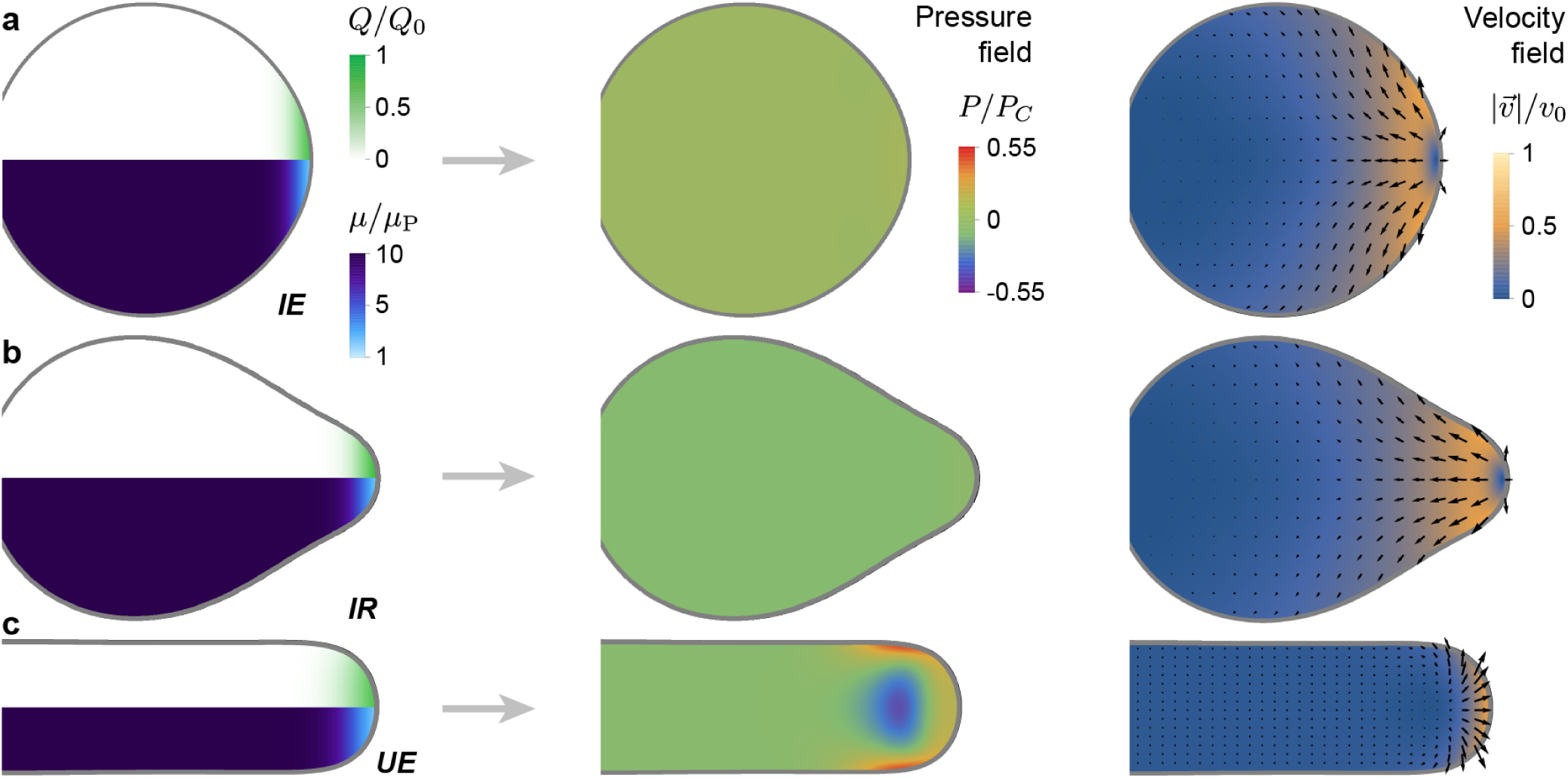
Distinct types of tissue dynamics during posterior body morphogenesis. **a-c**, Tissue pressure and velocity field associated to morphogenetic flows for the three limiting regimes shown in Fig. 2a, namely isotropic expansion (*IE*; a), intermediate regime (*IR*; b) and unidirectional elongation (*UE (nv)*; c). The parameter values for each of these cases are those indicated in Fig. 2a for each regime. The pressure and velocity fields during unidirectional tissue elongation are qualitatively different from those obtained during isotropic expansion and in the intermediate regime. The left column shows the input fields of the simulations, namely the tissue viscosity, *μ*(*x*), and the addition of new tissue material, *Q*(*x, y*), at the same time point as the pressure and velocity fields.

In contrast to isotropic expansion, in the regime where the tissue extends unidirectionally and posteriorly (Fig. 2a and Fig. 3b), most of the tissue material added at the posterior end flows posteriorly, causing the posterior tissue elongation. This is because it is less costly to create new tissue surface and extend the tissue posteriorly in order to accommodate the new material than moving the PSM material, as *σ*_A_ ≫ *σ*_C_ ∼ *σ*_P_. Just anterior of the region where new tissue material is added, the tissue starts flowing anteriorly and virtually arrests in the solid-like PSM. In this PSM region, the tissue is slightly compressed (positive, low pressure), indicating that posterior tissue elongation is enabled by MPZ tissues pushing on the solid-like PSM. The pressure profile displays a negative pressure zone in the medial region, just anterior of the region where new material is added, flanked mediolaterally by two regions of high pressure. This is because the tissue at the posterior-most end extends quickly posteriorly, whereas the solid-like anterior tissues cannot flow fast and follow it. The tissue in between the solid-like PSM and the posteriorly expanding MPZ needs to follow the posterior expansion at one end while maintaining connection with the PSM at the other, leading to an effective pull on the tissue and a negative pressure. It is important to note that this negative pressure region may be due to the 2D nature of these simulations, as in the full 3D geometry the capillary stresses from the tissue cross-section would create higher pressures in the tissue, likely preventing the formation of negative pressure regions. Yet, the reported spatial distribution of pressures would most likely remain qualitatively the same, with a medial region of small pressure localized anteriorly from where material is added. The high pressure regions flanking the low pressure medial region are due to the fact that the flow in this region encounters a very large anterior resistance due to the increasing viscosity towards the PSM and resistance to move mediolaterally due to tissue surface tension, effectively compressing the tissue.

These results indicate two limiting morphogenetic regimes, namely isotropic growth and unidirectional tissue expansion, which display both qualitative and quantitative distinct features.

### Morphogenetic flows during unidirectional tissue extension

In order to understand the types of morphogenetic flows that are involved in extending the posterior body axis, we explore the parameter space in the regime where there is a fluid-to-solid tissue transition (*μ*_A_ ≫ *μ*_P_) and the capillary stresses are less than or equal to the shear stress in the PSM, leading to unidirectional body elongation.

For large enough values of the capillary stress (*σ*_C_*/σ*_A_ *≥* 0.01) the system displays a source-type flow (Fig. 4a-c), with a single topological defect of topological charge +1 (Fig. 4b). As the capillary stresses are lowered or the PSM is made more rigid (*σ*_C_*/σ*_A_ ≲ 0.01; Fig. 4c), the system undergoes a sharp transition in the structure of the morphogenetic flow field, switching to a flow with three topological defects: two counter-rotating vortices symmetrically located about the midline (each with topological charge +1) and a stagnation point on the midline (with topological charge −1) (Fig. 4a,b). The total topological charge is conserved at the transition because the topology of the tissue boundary does not change. However, the number and spatial distribution of the topological defects change (Fig. 4b), leading to dramatic changes in the structure of the flow field (Fig. 4a). The counter-rotating vortices appear just anterior from the location where the tissue viscosity sharply increases (location of the fluid-to-solid tissue transition), and only for small capillary stresses. This is because in the limit of vanishing capillary stress, the new added material moving anteriorly eventually encounters the solid-like PSM and, since extending the tissue posteriorly has little cost in this limit, it progressively reverses its direction to medial, posterior-directed flows, generating the vortices. When the magnitude of tissue capillary stresses (tissue surface tension) is not negligible compared to the shear stresses in the system, deforming the tissue posteriorly has a finite cost and vortices are not observed.

**Figure 4.**
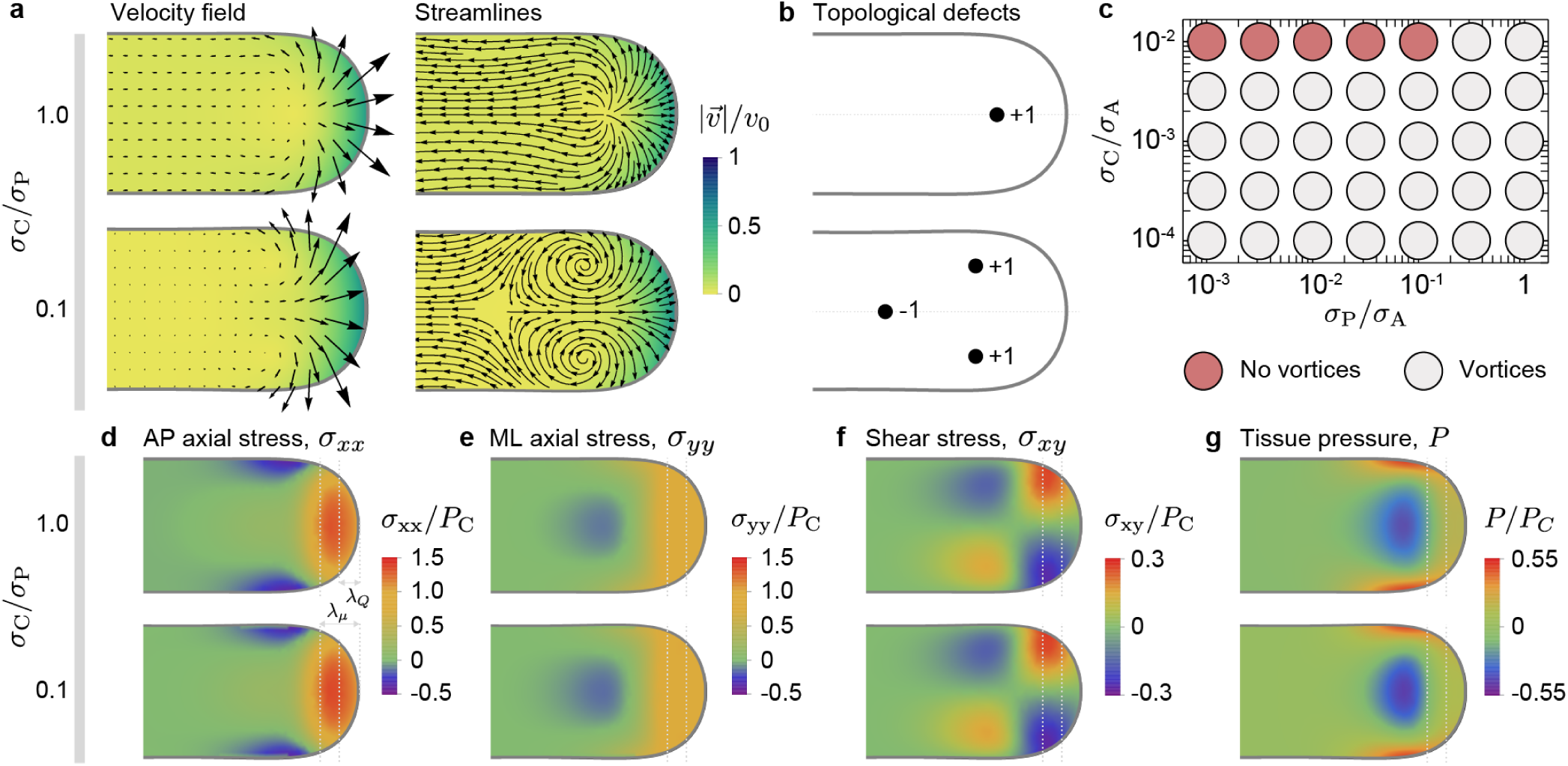
Morphogenetic flows and stress field during unidirectional elongation of the body axis. **a-b**, Velocity field and streamlines of morphogenetic flows for different values of *σ*_C_*/P*_*C*_, showing two structurally different flow fields (a). For large capillary stresses *σ*_C_, the flow emerges from a single topological defect (a; source) of charge +1 (b). For lower capillary stresses, the tissue flow dramatically changes its structure and is characterized by three topological defects, namely 2 counter-rotating vortices and a stagnation point. The remaining parameter values for each case are those indicated in Fig. 2a for unidirectional elongation (UE) in the absence and presence of vortices, labelled **nv** (no vortices) and **v** (vortices), respectively.**c**, Diagram indicating the structure (presence or absence of vortices) of tissue flows. The tissue morphogenetic flows sharply transit from the two flow structures shown in (a) as the parameters are changed. **d-g**, Spatial distribution of all components of the stress tensor, namely the AP axial stress *σ*_*xx*_ (d), the mediolateral stress *σ*_*yy*_ (e) and the shear stress *σ*_*xy*_ (f), as well as the tissue pressure *P* (g), for the two examples of tissue flows shown in panel (a). Despite the dramatically different structures of the flow field, the stresses are very similar. Both the location of the fluid-to-solid transition and the size of the region where cells enter ventral tissues are indicated by gray dashed lines at distances *λ*_*μ*_ and *λ*_*Q*_, respectively, from the posterior body end.

The transition in the structure of the flow field occurs with no qualitative changes in the stress field. In particular, the spatial distribution of all stress tensor components, namely the AP axial stress *σ*_*xx*_, the mediolateral (ML) stress *σ*_*yy*_ and the shear stress *σ*_*xy*_, in addition to the tissue pressure *P*, remain unchanged as the tissue flows change their structure dramatically at the transition (Fig. 4d-g). AP axial stresses *σ*_*xx*_ show large positive values in the fluid-like MPZ that push the tissue posteriorly against the tissue surface tension, which resists expansion. Far away from the posterior end, the anterior PSM is under AP compression because posterior tissues need to push on the PSM to extend the body posteriorly. In between these two limiting regions, as the tissue transitions from fluid-like to solid-like behavior, there is a medial region with low tensile stresses flanked by two lateral regions with large AP compressive stresses. The mediolateral stresses *σ*_*yy*_ display axial extension in the MPZ and compression in a medial PSM region anterior of the fluid-to-solid transition. The shear stress displays two high shear stress regions as the new tissue material in the MPZ flows out of the region, and it vanishes in the anterior PSM. Finally, the tissue pressure shows the characteristics described above for unidirectional posterior tissue elongation (Fig. 3c).

### Morphogenetic flows and tissue shape during posterior body axis elongation in zebrafish

In order to study the structure of morphogenetic flows and tissue shape during body axis elongation, we track cell movements in 3D throughout the tissue by following their nuclei (Fig. 5a; Supplementary Movie 1; Methods). To obtain the morphogenetic flows in ventral tissues, we defined a thin DV section through ventral tissues, coarse-grained the system by averaging the local cell velocities and calculated a dorsal projection (Methods). The velocity field and streamlines associated to the morphogenetic flows in the tissue display two counter-rotating vortices as the tissue transits from fluid to solid states (Fig. 5b; Supplementary Movie 2). Small tissue velocities are observed in the PSM, consistent with its solid-like state as also with previous observations^7^. In the fluid-like MPZ, higher velocities are observed in the outer part of the vortices, and slightly smaller velocities in the posterior-most, medial part of the MPZ, indicating that the tissue material (ingressing cells) entering from DM tissues tends to flow mediolaterally as they reach ventral tissues and incorporated in the vortices, with a smaller portion of this material directly moving posteriorly to elongate the body axis. The predicted structure of the morphogenetic flows from simulations (Fig. 5b) is remarkably similar to the experimentally measured structure of the morphogenetic flows, including its topological defects. Our simulations predict the observed counter-rotating vortices, as well as a stagnation point on the midline. The stagnation point is difficult to observe experimentally because of the presence of the notochord, which was not included in the simulations and likely imposes different boundary conditions in this region. While the general characteristics of the predicted velocity field are also observed experimentally, including small velocities in the PSM and larger velocities in the MPZ, our simulations predict the largest velocity magnitude at the posterior-most body end, whereas maximal speeds are experimentally observed in the outer part of the vortices. We believe the discrepancy is due to comparing 2D simulations with an experimental 2D projection of the 3D tissue flow. Overall, most features of the morphogenetic flows predicted for low tissue surface tension are observed experimentally, suggesting that capillary stresses are irrelevant (negligible compared to other stresses) during posterior body elongation in zebrafish.

**Figure 5.**
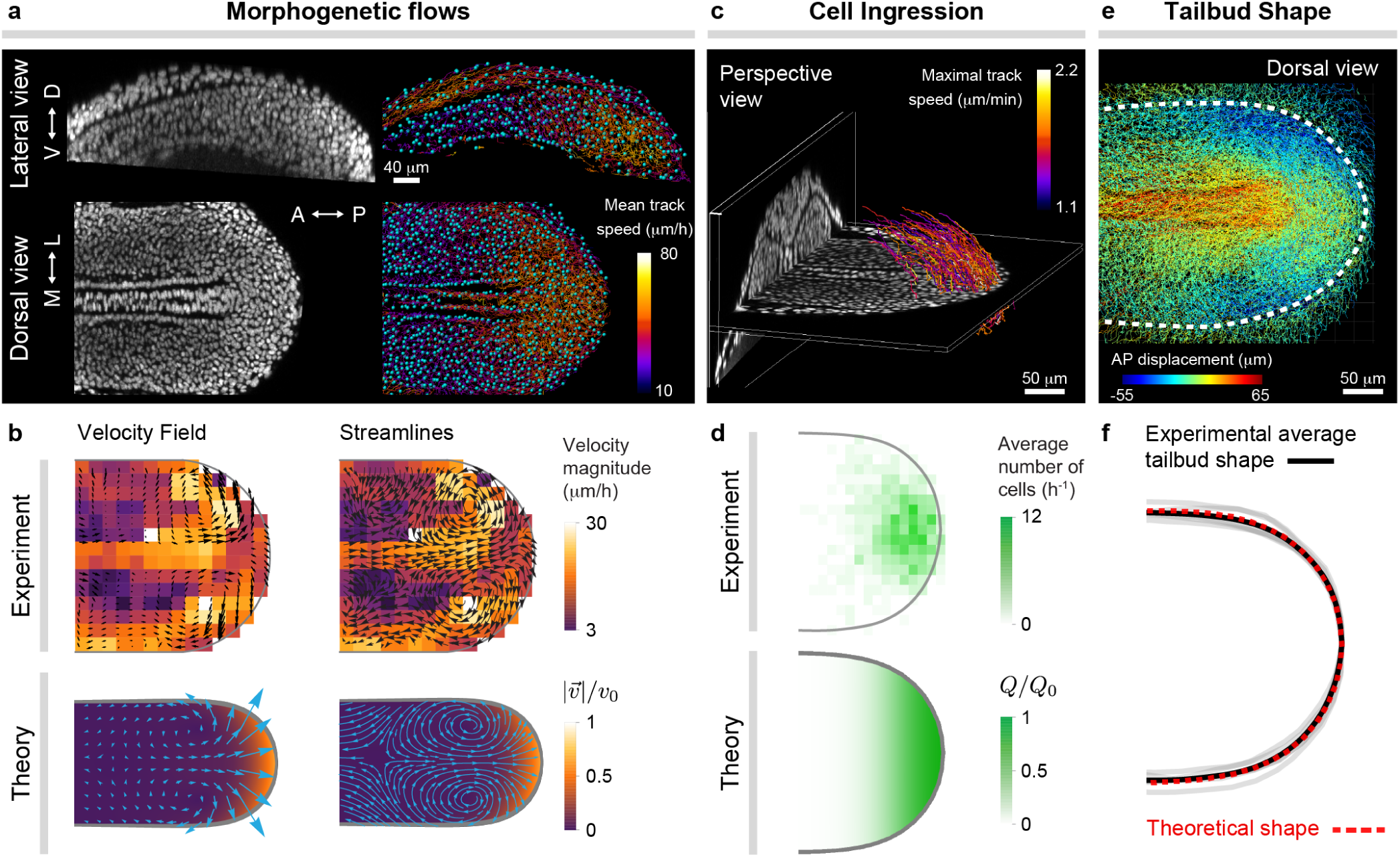
Morphogenetic flows and tissue shape during zebrafish posterior body axis elongation. **a**, Confocal lateral and dorsal sections of nuclei (gray) in posterior tissues of a zebrafish embryo at the 10 somite stage (left). Results of nuclear (cell) tracking showing a subset of identified nuclei (blue) and associated tracks (trajectories), color-coded according to their mean speed. **b**, Experimentally measured and theoretically obtained velocity field and streamlines of the morphogenetic flows in ventral tissues. **c**, Perspective 3D view of a confocal stack through elongating posterior tissues, showing a D-V section at the boundary between dorsal and ventral tissues and a AP section far from the posterior end. Tracks of cells entering ventral tissues from DM tissues are shown and color-coded according to their maximal speed. **d**, Measured frequency map of cells entering the MPZ from DM tissues and theoretically assumed spatial distribution of the same quantity, namely *Q*(*x, y*). **e**, Dorsal view of cell (nuclear) trajectories in posterior tissues, color-coded according to the magnitude of their AP displacement. The boundary of ventral tissues is shown as a thick dashed line and determines the shape of the elongating posterior body. **f**, Comparison of experimentally measured and theoretically predicted shapes of the posterior elongating tissue. The simulation results in panels b, d and f are all for the same parameters (*λ*_*μ*_ */λ*_*Q*_ = 1.5, *σ*_A_*/P*_C_ = 0.1, *σ*_P_*/P*_C_ = 0.02, *σ*_C_*/P*_C_ = 0.0001).

Beyond morphogenetic flows, tracking cellular movements in 3D allows a direct quantification of the spatial distribution of DM cells entering ventral tissues. We specifically tracked DM cells as they moved from DM tissues into ventral tissues (Fig. 5c; Methods) and obtained their frequency map (Fig. 5d; Methods), which should correspond to the theoretically defined rate of cell ingression from DM tissues into the MPZ, *Q*. Our assumed input function *Q* depends on the tissue pressure and is therefore also part of the solution of the simulations, even if the AP spatial extension is limited to a length scale *λ*_*Q*_ by construction (Fig. 5d). Experimentally, cells enter the MPZ through an approximately elliptical region adjacent to the posterior-most end of the body axis. While the input function *Q* and the experimentally measured frequency map of cells entering ventral tissues are slightly different, they share a key feature: cells enter ventral tissues within a limited distance (*λ*_*Q*_) from the posterior end. Indeed, the simulations reproduce the essential features of the experimental data despite these differences, suggesting the exact spatial distribution of *Q* is not essential as long as it is localized at the posterior-most end of the extending body axis.

One of the most important aspects of tissue morphogenesis is the tissue shape. In order to measure the shape of the tissue during posterior elongation, we also make use of the cell movements. Cell trajectories across the entire tissue seen from a dorsal view display distinctive AP displacements in the PSM and MPZ compared to surrounding tissues, allowing us to determine the boundary of these tissues (Fig. 5e; Methods). We quantified the 2D projected tissue shape and compared it to the tissue shape predicted in our simulations during unidirectional tissue elongation (and low tissue surface tension). The predicted tissue shape quantitatively reproduces the average shape of posterior body during axis elongation (Fig. 5f).

Overall, just considering the observed fluid-to-solid tissue transition along the AP axis and the observed flow of cells entering the MPZ from dorsal tissues in the simulations (with the same parameter set), it is possible to simultaneously reproduce the experimentally observed morphogenetic flows and tissue shape of the extending body axis.

## Discussion

We have studied the role of a spatially-localized fluid-to-solid tissue transition in the control of tissue morphogenesis in the specific case of zebrafish body axis elongation. By using finite element simulations to physically describe tissue morphogenesis at supracellular scales, and comparing the simulation results to quantitative experimental data for zebrafish axis elongation, we showed that unidirectional body elongation, morphogenetic flows and tissue shape can all be explained solely by the presence of a fluid-to-solid tissue transition along the AP axis. These results highlight the relevance of fluid-to-solid tissue transitions in the control of morphogenetic processes during embryogenesis.

Previous theoretical models have described cell movements during vertebrate axis elongation, albeit for DM tissues rather than ventral tissues. These simulations could not describe morphogenesis because a fixed tissue geometry (boundary) was assumed. In contrast, our description focuses on ventral tissues (PSM and MPZ) and describes the tissue as a continuum material that can undergo large shape changes, enabling the simulation of morphogenesis as a free-boundary problem. This continuum approach has been used to theoretically study tissue morphogenesis in different systems^8, 10^. However, since the tissue material properties and their spatiotemporal variations were unknown, previous works assumed spatially homogenous material properties (either solid or fluid), limiting their predictive power. Our description takes advantage of the detailed mechanical information recently available for zebrafish body axis elongation^2^, and accounts for the spatial variations in the key physical fields, namely the tissue material properties (solid/fluid tissue states) and growth (addition of tissue material through cell ingression). While active stresses do in general contribute to morphogenesis, we omitted them here because it has been shown that they do not contribute to posterior axis elongation in zebrafish^2^. Using the previously measured input fields, our simulations quantitatively reproduce unidirectional axis elongation, tissue morphogenetic flows and tissue shape. These results emphasize the importance of accounting for the spatiotemporal variations in all relevant physical fields (tissue growth, material properties and active stresses) to understand morphogenesis^1^.

Our theoretical results indicate that in the absence of a spatially-localized fluid-to-solid tissue transition, posterior tissues expand isotropically or display considerable lateral spreading, but fail to form a body axis with sustained unidirectionally elongation at its posterior end. Moreover, large enough tissue tensions completely halt axis elongation. While several morphological phenotypes and even the ceasing of posterior elongation have been observed in zebrafish mutants or under specific molecular perturbations^7, 24–26^, it is unclear if these observed phenotypes are related to those predicted in our description, as there are no measurements of physical parameters in those experimental conditions. In contrast, for low tissue surface tension and in the presence of a fluid-to-solid transition in the tissue physical state along the AP axis our simulations predict unidirectional axis elongation. These results indicate that the observed progressive rigidification of posterior tissues^2^, as cells in the fluid-like MPZ get incorporated into the solid-like PSM, is necessary to explain the sustained posterior elongation of the body axis. More generally, it is possible that a progressive and regionally-controlled tissue rigidification may be necessary to shape other embryonic structures, especially in cases that involve tissue elongation. Our work shows that regardless of the specific physical mechanism underlying the fluid-to-solid tissue transition (e.g., cellular jamming/glass transitions, extracellular matrix rigidification, etc.), the spatiotemporal control of physical (fluid-solid) state of the tissue is an important physical mechanism to control morphogenetic flows and tissue morphogenesis.

The predicted morphogenetic flows for simulated unidirectional body elongation, indicate the presence of a topological transition in the structure of morphogenetic flows, with two distinct flow patterns exist depending on the values of tissue viscosity and surface tension. In the presence of a fluid-to-solid tissue transition along the AP axis and for low enough tissue tensions, the simulated morphogenetic flows during unidirectional axis elongation display two counter-rotating vortices as the tissue transits from fluid to solid-like states, as well as an stagnation point in the PSM. Our experimental observations show that this predicted flow pattern, which arises naturally from the dynamics of the system, is observed in zebrafish ventral tissues during posterior body elongation (Fig. 5a). For large enough values of tissue surface tension, vortices are suppressed and tissue flows display a structure with a single source (defect). While not observed in wild type conditions, it may be possible that zebrafish mutants display these kind of flows. The structure of the morphogenetic flow and, especially a sharp transition between very different structures of the flow, may have important consequences for proper embryonic development. The signals that cells experience in the embryo depend on their physical trajectories, which may be completely different in the two distinct flow structures predicted, suggesting that the flow structure may play an important role in controlling cell behavior during embryogenesis.

Beyond tissue flows, the spatial distribution of the different stress components provides information about the physical mechanism of posterior body elongation. The new added tissue material in the fluid-like MPZ provides the necessary pushing force to extend the body axis, overcoming the resistance from a potential tissue surface tension (if present). This highlights an important mechanical role of dorsal tissues in posterior elongation, as the tissue forces in dorsal tissues that drive cell insertion into the MPZ translate to large AP stresses in the MPZ that enable body elongation. However, posterior tissues need to push on something to elongate unidirectionally. The reported compressive AP stresses in the anterior PSM indicate that posterior tissues push on the PSM to sustain posterior elongation. While the posterior MPZ tissue needs to be in a fluid state to enable tissue remodeling (cells entering from DM tissues) and morphogenesis, tissue rigidification in the PSM is necessary to mechanically support the posterior extension of the body axis.

Our simulations quantitatively reproduce the experimentally observed unidirectional elongation of the body axis, the structure of tissue flows (with vortices) and the tissue shape at the extending posterior end, for values of the capillary stress (tissue surface tension) much smaller than the shear stresses in the tissue, suggesting that the tissue surface tension is not important to understand posterior elongation *in vivo*. Measurements of tissue surface tension during body axis elongation are not available, but previously measured values of tensions in epithelial tissues are consistent with this result^22, 23^. The mechanics of the tissue surface may, however, be more complex. While previous observations have shown that no extracellular matrix is present between cells in the PSM and MPZ tissues^27^, there is a fibronectin sheath at the tissue boundary that could change our description of the tissue surface mechanics and potentially affect tissue elongation. However, disruption of the fibronectin sheath does not seem to prevent the formation of the body axis at the developmental stage studied here^27–29^. While spatiotemporal variations in the extracellular matrix sheath can also partially contribute or facilitate posterior extension, these observations, our simulations and recent measurements of tissue mechanics during axis elongation indicate that rigidification of the PSM (i.e., the fluid-to-solid transition) helps to mechanically sustain the posterior extension of the body axis.

Altogether, our results indicate that the presence of a fluid-to-solid transition in the tissue physical state along the AP axis, combined with the posterior addition of new tissue material, is sufficient to simultaneously explain the observed unidirectional posterior elongation of the body axis, the morphogenetic flows in the tissue and the shape of the elongating axis. These results highlight the important role of transitions in the tissue physical state for the control of embryonic morphogenesis. Future studies will determine whether this physical mechanism of morphogenesis guides the formation of other embryonic structures.

## Methods

### Computational Methods

We solve Eqs. 3-5 with the boundary conditions described above using Comsol Multiphysics 5.3, which employs Finite Element Methods. Laminar flow and moving mesh models were used to simulate large deformations of the tissue material under growth. The Comsol model consists of a box, with sides of length 50 times larger than the tissue size, filled with a fluid of negligible viscosity, and the tissue initially contained in a semicircular region with both ends fixed to one side of the box. The tissue satisfies the source and viscosity profiles (Eqs. 1, 2). Fluid addition at the tip and the isotropic term (∇· **u**) in Eq. 6 are directly inputted into the weak solution integration formulation of Comsol. No tissue flow can go through the box and pressure is set to zero at far away walls. The system is meshed with the smallest element being 130 times smaller than the radius of the initial semicircular area close to the tissue-fluid and tissue-box interfaces, with the mesh updating as needed to accommodate large deformations resulting from tissue growth.

### Zebrafish husbandry, lines and experimental manipulations

Zebrafish (*Danio rerio*) were maintained under standard conditions^30^. Animal husbandry and experiments were done according to protocols approved by the Institutional Animal Care and Use Committee (IACUC) at the University of California, Santa Barbara. Nuclei were labeled to track cell movements by either using the Tg(h2afva:GFP)^*kca*6^ transgenic line^31^ or by injection with 80-100pg H2B-RFP mRNA at 1-2 cell stage.

### Embryo imaging

In all experiments, 8-10 somite stage zebrafish embryos were mounted in 1% low-melting point agarose in a glass bottom petri dish (MatTek Corporation) for a dorsal view of the tailbud and imaged at 25*°*C using an inverted Zeiss Laser Scanning Confocal (LSM 710, Carl Zeiss Inc.). Confocal stacks through the tailbud were acquired with a step size of 2 *μm* and time interval of 2 minutes for 2 hours, using a 25x water immersion objective (LD LCI Plan-Apochromat 25x/0.8 Imm Corr DIC M27, Carl Zeiss Inc.).

### Cell movement tracking

Imaging data was first processed using Imaris (Bitplane). Obtained confocal stacks through posterior tissues were first smoothed using a 1-pixel Gaussian filter. These stacks were then corrected for photo-bleaching using the normalize timepoints function. Finally, attenuation correction was applied to correct for z-attenuation. If required, the free-rotate tool was used to align the data such that all samples had the same alignment with respect to a specified Cartesian reference frame. The Measurement Points tool was used to identify the location of the notochord and tissue boundaries as well as to define the plane of cell ingression. After processing, nuclei were detected using the spots function, and tracked using the Brownian motion algorithm. Nuclei positions and velocities were output for further analysis.

### Cell ingression analysis

To quantify cell ingression rate into ventral tissues from DM tissue, we selected cells that exhibited displacements in the dorsal-ventral axis larger than 30*μm*. For each selected cell trajectory, we determined all time points that the cell is located in the dorsal-ventral boundary region (10*μm* thickness) and then calculated the average x and y components of the cell position over the identified time points for which the cell is transiting from dorsal to ventral tissues. Cell ingression positions are measured over 5 distinct samples. To binning cell ingression rate from different samples, body axes are rescaled by the distance between the posterior end of the body and posterior end of notochord. Cell ingression rate is binned in terms of x and y positions.

### Velocity field analysis

Nuclei trajectories were determined using the particle tracking algorithm in Imaris and subsequently used to compute 2D velocity field. For each cell trajectory in 3D, a B-spline curve is computed to eliminate high frequency noise. Cell velocities are then numerically computed from the B-spline curve. The obtained velocity values are averaged spatially and temporally (coarse-graining) to obtain a smooth velocity field. The coarse-grained velocity field is then projected on the xy plane (dorsal view) and binned in terms of x and y positions with a bin width 10*μm*.

## Acknowledgments

We thank all members of the Campàs group for their comments and help. OC thanks Mark Bowick (KITP, University of California, Santa Barbara) for insightful comments. This work was supported by the Eunice Kennedy Shriver National Institute of Child Health and Human Development of the National Institutes of Health (R01HD095797).

## Author contributions statement

SPB, EKC, PR and OC designed research; OC supervised the project; SPB, EKC and PR performed simulations of tissue dynamics; GS-V performed the experiments; SK, GS-V and SPB analyzed data; SPB, PR and OC wrote the manuscript. All authors reviewed the manuscript.

